# How the window of visibility varies around polar angle

**DOI:** 10.1101/2024.07.12.603257

**Authors:** Yuna Kwak, Zhong-Lin Lu, Marisa Carrasco

**Affiliations:** Department of Psychology, New York University, New York, United States; Department of Arts & Sciences, New York University Shanghai, Shanghai, China; Center for Neural Science, New York University, New York, United States

## Abstract

Contrast sensitivity, the amount of contrast required to detect or discriminate an object, depends on spatial frequency (SF): The Contrast Sensitivity Function (CSF) peaks at intermediate SFs and drops at lower and higher SFs and is the basis of computational models of visual object recognition. The CSF varies from foveal to peripheral vision, but only a couple studies have assessed changes around polar angle of the visual field. Sensitivity is generally better along the horizontal than the vertical meridian, and better at the lower vertical than the upper vertical meridian, yielding polar angle asymmetries. Here, we investigate CSF attributes at polar angle locations at both group and individual levels, using Hierarchical Bayesian Modeling. This method enables precise estimation of CSF parameters by decomposing the variability of the dataset into multiple levels and analyzing covariance across observers. At the group level, peak contrast sensitivity and corresponding spatial frequency with the highest sensitivity are higher at the horizontal than vertical meridian, and at the lower than upper vertical meridian. At an individual level, CSF attributes (e.g., maximum sensitivity, the most preferred SF) across locations are highly correlated, indicating that although the CSFs differ across locations, the CSF at one location is predictive of the CSF at another location. Within each location, the CSF attributes co-vary, indicating that CSFs across individuals vary in a consistent manner (e.g., as maximum sensitivity increases, wso does the SF at which sensitivity peaks), but more so at the horizontal than the vertical meridian locations. These results show similarities and uncover some critical polar angle differences across locations and individuals, suggesting that the CSF should not be generalized across iso-eccentric locations around the visual field. Our window of visibility varies with polar angle: It is enhanced and more consistent at the horizontal meridian.

**Author summary:** The contrast sensitivity function (CSF), depicting how our ability to perceive contrast depends on spatial frequency, characterizes our “window of visibility”: We can only see objects with contrast and spatial frequency properties encompassed by this function. The CSF is mostly assessed only along the horizontal meridian of the visual field and sometimes averaged across locations, but visual performance varies with polar angle (e.g., we are more sensitive to objects along the horizontal than the vertical meridian). Here, we systematically assess the key attributes of the CSF and show critical differences in the window of visibility across polar angles and individuals. We found that at the horizontal meridian, our overall contrast sensitivity and preferred SF are higher, and CSFs of individual observers co-vary more than at the vertical meridian. This research highlights that this fundamental perceptual measure is not the same and should be assessed around the visual field. Polar angle thus should be a key consideration for applications of the CSF in computational models of vision.

## Introduction

*Contrast sensitivity*, the ability to discriminate visual patterns from a uniform background, is closely related to performance in daily tasks, such as face recognition (Wolffsohn, Eperjesi, & Napper, 2005), reading (Brussee, van den Berg, van Nispen, & van Rens, 2017; Rubin & Legge, 1989), mobility (Marron & Bailey, 1982), and driving (Owsley & McGwin, 2010; Schieber, Kline, Kline, & Fozard, 1992). It is also central for clinical assessment (Owsley, 2003; Pelli & Bex, 2013), as contrast sensitivity is impaired in ophthalmic conditions such as myopia (Anders et al., 2023; Baldwin et al., 2023; Collins & Carney, 1990; Ginsburg, 2006; but see Xu et al., 2022) and amblyopia (Hou et al., 2010; Levi & Li, 2009; Zhou et al., 2006), and in neurological conditions such as cerebral lesions (Bodis-Wollner, 1972) and schizophrenia (Cimmer et al., 2006).

Contrast sensitivity varies substantially as a function of spatial frequency (SF): The *contrast sensitivity function (CSF)* characterizes an individual’s ability to reliably discriminate different levels of SF at a location, and is typically bandpass, peaking at mid SFs (**Figure 1a**). It is considered the window of visibility because objects with properties within the CSF-specified region are visible (Campbell & Robson, 1968). The CSF has also served as the front-end filter in observer models of object recognition, in which the CSF controls the gain for different spatial frequency components of objects (Chung, Legge, & Tjan, 2002; Hou, Lu, & Huang, 2014; Majaj, Pelli, Kurshan, & Palomares, 2002; Rovamo, Luntinen, & Näsänen, 1993; Watson & Ahumada, 2005). Because the CSF provides critical information about spatial vision, efforts have been made to model how it varies with different stimulus conditions (Graham, 1989; Watson, 2018), including luminance (e.g., Rovamo, Virsu, & Näsänen, 1978), size (e.g., Rovamo et al., 1993), stimulus duration (e.g., Legge, 1978), temporal characteristics (e.g., Kelly, 1979; Robson, 1966), and eccentricity (e.g., Hilz & Cavonius, 1974; Rovamo et al., 1978). For instance, at farther eccentricities, the CSF shifts downward and toward lower SFs, reducing the window of visibility (e.g., Virsu & Rovamo, 1979).

**Figure 1.**
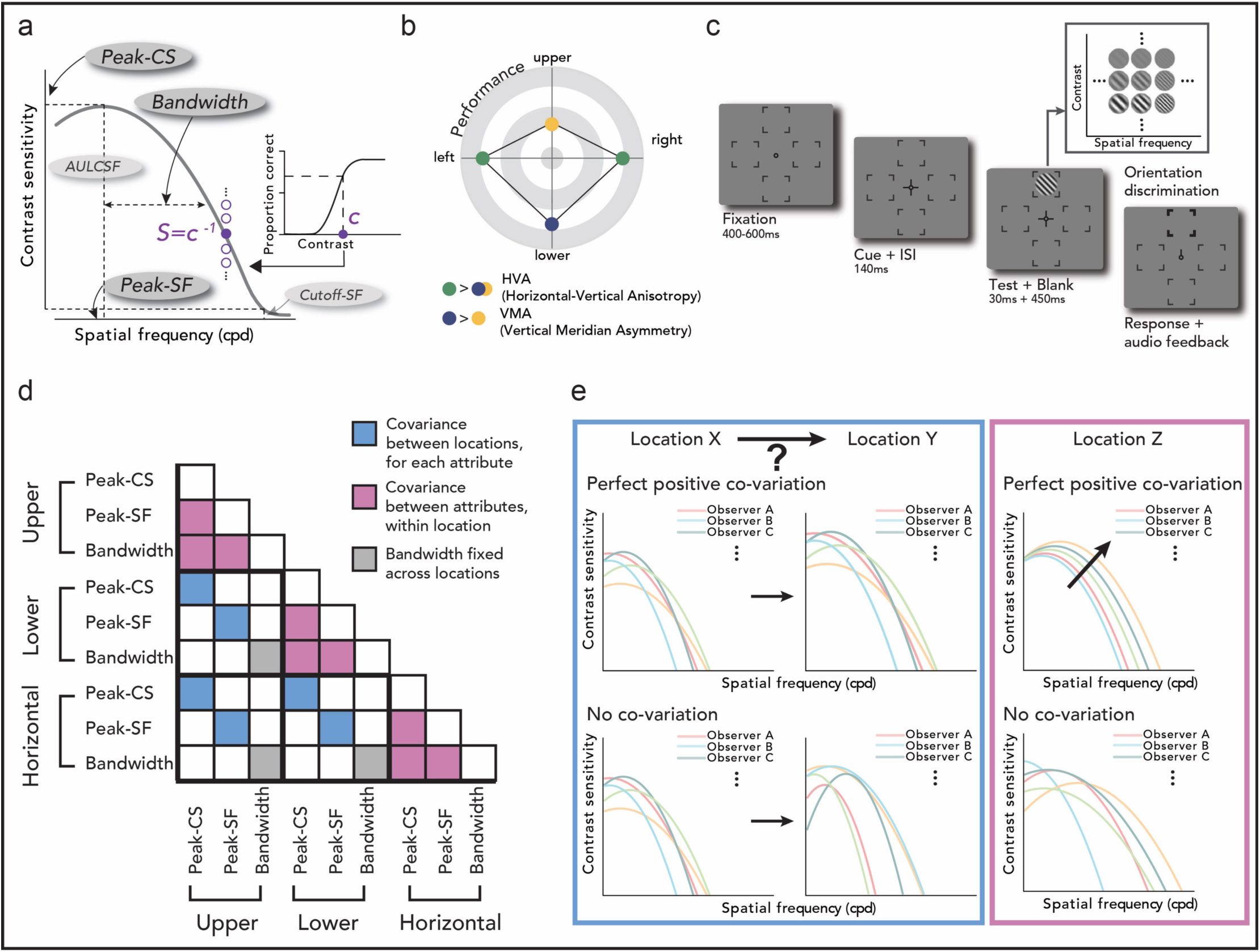
Experimental schematics and procedure. **(a) CSF (Contrast Sensitivity Function) analysis.** The CSF depicts how contrast sensitivity depends on spatial frequency (SF). The inverse of the contrast threshold (c^−1^) is contrast sensitivity (S) at a particular SF. Bayesian inference was used to estimate the CSF parameters —peak contrast sensitivity (peak-CS: maximum sensitivity), peak spatial frequency (peak-SF: the most preferred SF), and bandwidth (closely related with CSF shape). Bandwidth was fixed across locations (see text and **Methods**), and results of the cutoff spatial frequency (cutoff-SF: the highest discernable SF) and area under the log CSF (AULCSF: total sensitivity) are reported in **Supporting Information**. Higher values of these attributes are related to higher sensitivity, better performance, and/or a wider range of visible SFs and contrasts. **(b) Polar angle asymmetries.** In many visual dimensions, performance is higher at the horizontal than the vertical (HVA), and at the lower than the upper vertical meridian (VMA). **(c) Task.** A test Gabor stimulus was presented briefly at one of the four polar angle locations, and participants judged its orientation (CW or CCW). SF and contrast of the stimulus varied on each trial. **(d) Schematics of covariance analysis.** The covariance of the three CSF attributes (peak-CS, peak-SF, bandwidth) for all three locations is a 9 x 9 matrix. The covariation of locations for each attribute (blue cells), and the covariation of attributes within each location (pink cells) were assessed. Covariance for bandwidth is shaded in gray because the bandwidth was identical across locations for each observer in the HBM and thus, the covariance is not informative in most cases (but see Figure 4). **(e) Interpretation of covariance analysis.** Covariance of locations for each attribute (**d**, blue cells) tests whether CSFs and their corresponding key attributes co-vary as a function of polar angle. For example, a perfect positive correlation (coefficient of 1) for peak-CS among locations X and Y, indicates that an observer with a higher peak-CS at location X than other individuals also has a higher peak-CS at location Y. (**e**, blue panel top row). This relation does not hold for a scenario where there is no correlation (coefficient of 0) for any CSF attributes between locations X and Y. Covariance within each location (**d**, pink cells) allows investigation of how individuals’ CSFs relate to one another within each location. A perfect positive covariation (correlation coefficient of 1) of all combinations of attributes within a location indicates that observers’ CSFs are organized diagonally in the SF-contrast space: the higher the peak-CS, the higher the peak-SF, and the wider the bandwidth (**e**, pink panel, top row). The fewer the correlations, the more variability in the pattern of individual differences (**e**, pink panel, bottom row is a scenario where no correlations are present among CSF attributes within a location).

Most studies in vision have assessed visual performance only along the horizontal meridian, but visual information in the real world is distributed around the entire visual field. Behavioral sensitivity in many visual dimensions varies around polar angle (**Figure 1b**; review, Himmelberg et al., 2023): When eccentricity is constant, contrast sensitivity at a particular SF is better along the horizontal than the vertical meridian (HVA: Horizontal-Vertical Anisotropy), and better at the lower than the upper vertical meridian (VMA: Vertical Meridian Asymmetry), indicating polar angle asymmetries (Abrams, Nizam, & Carrasco, 2012; Baldwin, Meese, & Baker, 2012; Carrasco, Talgar, & Cameron, 2001; Himmelberg, Winawer, & Carrasco, 2020). Moreover, cortical surface area (Benson, Kupers, Barbot, Carrasco, & Winawer, 2021; Himmelberg et al., 2021; Himmelberg, Winawer, & Carrasco, 2022), and the distribution of cone photoreceptors and retinal ganglion cells (Curcio & Allen, 1990; Webb & Kaas, 1976) vary as a function of polar angle. These differences indicate that measurements along the horizontal meridian do not generalize to other locations. However, scarce attention has been paid to characterizing the CSF at iso-eccentric locations around the visual field. There are only two studies that systematically assessed the entire CSF at polar angle meridians and showed that the these asymmetries are present for contrast sensitivity across a wide range of SFs (Jigo, Tavdy, Himmelberg, & Carrasco, 2023; Kwak, Zhao, Lu, Hanning, & Carrasco, 2024). Interestingly, M scaling - scaling the stimulus to equate its V1 cortical representation at different locations (Rovamo & Virsu, 1979) - eliminates eccentricity, but not polar angle differences in CSF (Jigo et al., 2023). In addition, behavioral asymmetries around polar angle are intensified at higher SFs (Cameron, Tai, & Carrasco, 2002; Carrasco et al., 2001; Himmelberg et al., 2020; McAnany & Levine, 2007), underscoring the need to characterize the entire CSF. Here, we conducted a systematic evaluation of the CSF around polar angle by investigating key attributes at each location (**Figure 1a**). Importantly, for the first time, we examine the relations in CSFs across locations and individual observers.

Given the prominence of the CSF as a building block of spatial vision, obtaining accurate CSF estimates is essential (Glassman et al., 2024; Hou et al., 2010; Lesmes, Lu, Baek, & Albright, 2010). We extracted key CSF attributes at the upper vertical, lower vertical, and horizontal meridian (**Figure 1a,c**), using Hierarchical Bayesian Modeling (HBM; **Methods** and **Figure S1**), which enables accurate and precise CSF estimates with relatively few observations (Zhao, Lesmes, Hou, & Lu, 2021): Peak-CS is the maximum contrast sensitivity, peak-SF is the SF corresponding to the peak-CS and thus the most preferred SF, and bandwidth is the width of the CSF (cutoff-SF -the highest perceivable SF- and AULCSF -area under the log CSF, the total sensitivity- are in **Supporting Information**) We found that across polar angle locations, the shape of the CSF is constant, but that sensitivity (e.g., peak-CS and peak-SF) is higher at the horizontal than the vertical (HVA), and at the lower than the upper vertical meridian (VMA), consistent with the two studies from our group that have measured CSF attributes around polar angle (Jigo et al., 2023; Kwak et al., 2024).

Importantly, to examine the relations in CSFs across locations and across individual observers, we analyzed the covariance of the CSF attributes (**Figures 1d-e**). First, across locations, CSF attributes were correlated (e.g., peak-CS at horizontal and peak-CS at upper vertical meridian), indicating that an observer with an enhanced CSF (e.g., higher contrast sensitivity) at one location also had enhanced CSFs at other locations, compared to other observers (for a possible scenario, see **Figure 1e** blue panel top row). Second, within each location, CSF attributes were highly correlated with one another (e.g., peak-CS at horizontal and peak-SF at horizontal meridian), indicating that individuals’ CSFs differ in a systematic manner (for a possible scenario, see **Figure 1e** pink panel top row), especially at the horizontal meridian. Observers’ CSFs are shifted along the diagonal direction in the log(SF)-log(contrast sensitivity) space. The study reveals critical differences in and relations among CSFs around polar angle, at both group and individual levels, indicating that CSFs should not be generalized around the meridians.

## Results

We estimated CSF parameters - peak contrast sensitivity (peak-CS: maximum sensitivity), peak spatial frequency (peak-SF: the most preferred SF), and bandwidth (full width at half maximum sensitivity, related to CSF shape) - using Bayesian inference (**Figure 1a**). The likelihood of the observed data was computed by fitting psychometric functions with contrast thresholds *(c* = S^−1^) set as the inverse of the contrast sensitivity (S) defined by the parameters, and Bayes’ Rule was used to infer the posterior distribution of the CSF parameters. Instead of fitting the CSF model separately for each participant and location, we used a Hierarchical Bayesian Model (HBM) which considers potential relations among parameters across the hierarchy, to accurately estimate CSF parameters (**Methods** and **Figure S1**). In addition, cutoff spatial frequency (cutoff-SF: the highest perceivable SF, or SF at S = 1) and the area under the log CSF (AULCSF: total window of visibility) were extracted from the fitted CSFs and are reported in **Supporting Information**. Higher values of these attributes indicate higher sensitivity, better performance, and/or a wider range of visible SFs and contrasts.

The HBM were fit to the data from the upper vertical, lower vertical, and the horizontal meridians. Data were collapsed across the left and right horizontal locations, as previous studies have shown that contrast sensitivity is similar across these two locations (e.g., Barbot et al., 2021; Cameron et al., 2002; Jigo et al., 2023).

### CSF shape is the same across polar angle locations

It is well established that Peak-CS and Peak-SF vary as a function of eccentricity (e.g., Virsu & Rovamo, 1979), but less is known regarding bandwidth, which determines the shape of the CSF. To test whether the CSF shape differs around polar angle, we fitted two versions of the HBM to the data (**Methods**) - one in which bandwidth was constrained to be the same across the three locations for each observer, and the other in which no such constraint was imposed. The BPIC (Bayesian predictive information criterion; Ando, 2007) values used to quantify the goodness of fit was 22,141 for the fixed and 22,195 for the varying model (lower values indicate better fit). The fixed model provided statistically equivalent fits as the most saturated model, suggesting that - for each individual, the shape of the CSF does not differ around polar angle locations, and CSFs around polar angle are translationally invariant in the log(SF)-log(contrast sensitivity) space. We used the results from the fixed bandwidth model for further analyses (mean bandwidth across observers = 2.908 octaves) and report our findings focusing on peak-CS and peak-SF in the following sections (but see **Figure 4**).

### CSFs are enhanced at the horizontal meridian compared to the vertical meridian

To qualitatively preview the results, there were contrast sensitivity differences around polar angle at the group level (**Figure 2a**): Contrast sensitivity across SFs was higher for the horizontal than the vertical meridian (HVA:

**Figure 2.**
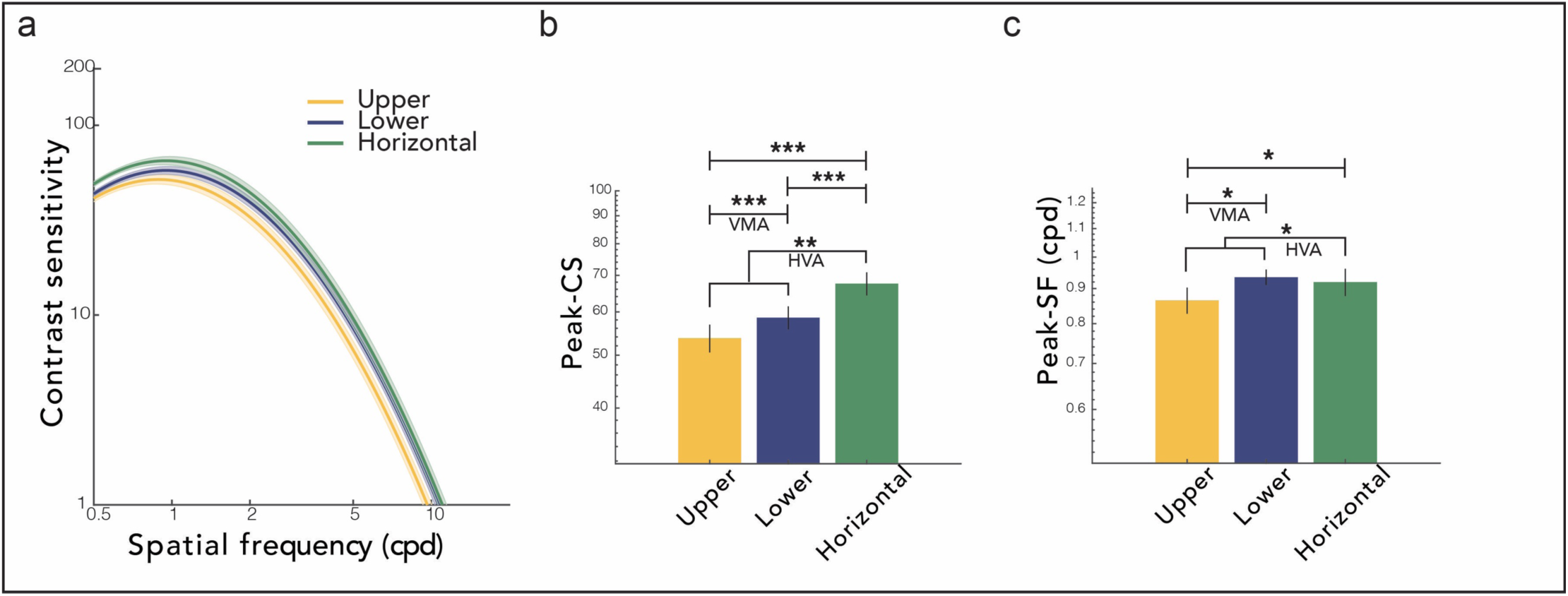
Polar angle sensitivity differences. **(a) CSFs.** An example participant’s data are shown in **Figure S2**. **(b) Peak contrast sensitivity. (c) Peak spatial frequency.** Results for cutoff-SF and AULCSF are shown in **Figure S3**. \*\*\**p* < 0.001, \*\**p* < 0.01, \**p* < 0.05. Error bars are ±1SEM. HVA and VMA denote Horizontal-Vertical Anisotropy and Vertical Meridian Asymmetries, respectively.

Horizontal-Vertical Anisotropy), and for the lower than the upper vertical meridian (VMA: Vertical Meridian Asymmetry) (for an example participant’s trial-by-trial data and the best fitting CSF models for each location, see **Figure S2**). We examined each of the key CSF attributes for a detailed understanding of this pattern.

We conducted one-way repeated measures ANOVA, with location (upper, lower, horizontal) as a within-subject factor, and observed significant location effects for peak-CS (*F*(2,27) = 46.292, *p* < 0.001, n^2^ = 0.632) and peak-SF (*F*(2,27) = 4.166, *p* = 0.025, n^2^ = 0.134).

For peak-CS (**Figure 2b**), there were clear HVA and VMA. Peak-CS was significantly higher at the horizontal than the vertical meridian (average of upper and lower), indicating HVA (*t*(27) = 3.318, *p* = 0.003, *d* = 0.754). It was also higher at the lower vertical than the upper vertical meridian, indicating VMA (*t*(27) = 3.318, *p* < 0.001, *d* = 0.333). In addition, the peak-CS at the horizontal was higher than at the lower vertical meridian (*t*(27) = 6.001, *p* < 0.001, *d* = 0.591).

For peak-SF (**Figure 2c**), we observed both HVA and VMA. The CSF peaked at a higher SF at the horizontal than at the vertical (*t*(27) = 2.530, *p* = 0.018, *d* = 0.102), and at the lower vertical than the upper vertical meridian (*t*(27) = 2.530, *p* = 0.026, *d* = 0.387). There was no difference between the horizontal and the lower vertical meridian.

The cutoff-SF and AULCSF also differed significantly as a function of polar angle, in a direction consistent with peak-CS and peak-SF (**Figure S3)**. These results indicate that the CSF and its corresponding attributes vary around the visual field and should not be generalized across locations.

### Individual CSFs across locations are highly correlated

Sensitivity at the group level differed around polar angle (**Figure 2**), indicating that the CSF and its attributes did not generalize across iso-eccentric locations around the visual field. But are *individual* CSFs at these locations related to each other, such that for example, an observer with a higher peak-CS than other observers at one location also have a higher peak-CS at another location?

To answer this question, we analyzed subsections of the covariance matrix (**Figure 1d** blue cells) and examined how each CSF attribute of an observer co-varies *across* locations. If all CSF attributes perfectly co-vary across pairs of locations (**Figure 1e** blue panel top row), observers’ CSF would be shifted and scaled by the same amount between the two locations, preserving individual variability across locations. If none of the CSF attributes co-vary across locations (**Figure 1e** blue panel bottom row), individual variability would not be preserved across locations.

We found that the peak-CS at one location was positively correlated with their counterparts at all other locations (**Figure 3a**). The same pattern was observed for peak-SF (**Figure 3b**). Therefore, CSF attributes strongly co-vary across all polar angle locations (more consistent with **Figure 1e** blue panel top row, than with the bottom row). These findings indicate that individual variability is preserved across all polar angle locations and suggest that an observer’s CSF and its attributes at one location are good predictors of those at another location. For the respective correlations for cutoff-SF and AULCSF, see **Figure S4**.

**Figure 3.**
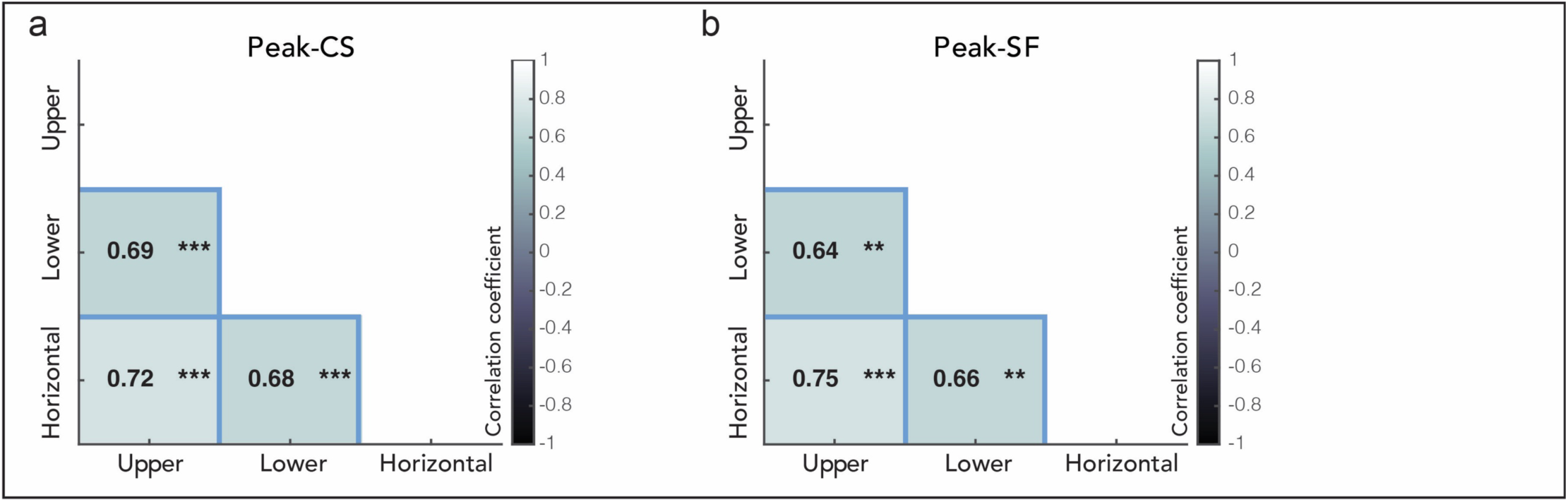
Covariation of each CSF attribute across polar angle locations. **(a) Peak contrast sensitivity. (b) Peak spatial frequency.** Each cell corresponds to a blue cell in Figure 1d. The covariation of cutoff-SF and AULCSF are shown in **Figure S4**, and confidence intervals are in **Figure S7**. For observers’ data examples, see **Figure S5**. \*\*\**p* < 0.001, \*\**p* < 0.01.

### CSFs attributes are correlated within each location, and more so for the horizontal meridian

Next, we assessed the relation among CSF attributes *within* each location at the individual level (**Figure 4**). We asked: How similar or different are individual CSFs within each location, and does this extent differ around polar angle? We analyzed triangular sections below the main diagonal in the covariance matrix (**Figure 1d** pink cells) to investigate whether and how key CSF attributes co-vary with another, across observers, separately for each location. Note that here, the co-variation of bandwidth with other CSF attributes is informative because the bandwidth was free to vary across observers (**Methods**; Note that in previous sections focusing on comparisons across locations, the bandwidth is not informative as it was fixed across locations). The more significant correlations across pairs of CSF attributes at a location (both positive and negative), the more consistently individual CSFs at that location vary in the log(SF)-log(contrast sensitivity) space (**Figure 1e** pink panel top row). The fewer the correlations among CSF attributes, the more variability among individual CSFs (**Figure 1e** pink panel bottom row).

**Figure 4.**
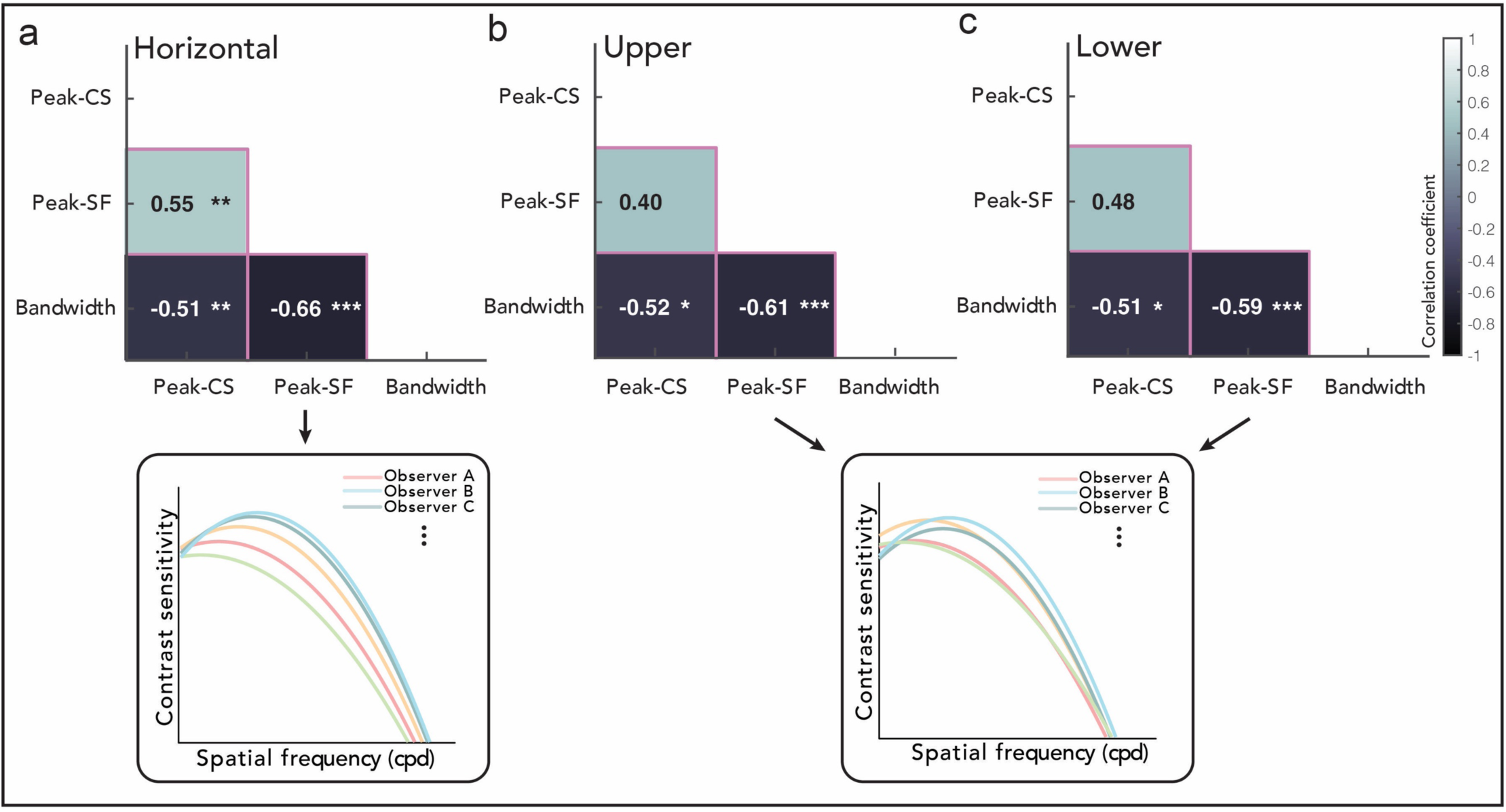
Covariance of CSF attributes within each location (pink cells in Figure 1d-e). Correlation between pairs of attributes at the **(a)** Horizontal, **(b)** Upper vertical, and **(c)** Lower vertical meridian. Note that here, the bandwidth is informative because the bandwidth varies as a function of individual observer (it is fixed only across locations). The color of each cell corresponds to the strength of correlation. Inset demonstrates schematics of simulated individual CSFs at each location, assuming that the significant/non-significant correlations are perfect/null correlations (coefficient of 1/0) for simplicity. Each cell corresponds to a pink cell in Figure 1d. The covariation of all attributes including cutoff-SF and AULCSF are shown in **Figure S6**. Confidence intervals are in **Figure S7**. \*\*\**p* < 0.001, \*\**p* < 0.01, \**p* < 0.05. Black and white letters are for visibility.

The extent to which the CSF attributes correlations differ around the polar angle is depicted in **Figure 4** (see **Figure S5** for examples of individual data). Only at the horizontal meridian, *all* CSF attributes co-vary with one another (**Figure 4a**). The peak-CS and peak-SF are positively correlated, indicating a shift, across observers, along the diagonal direction in the log(SF)-log(contrast sensitivity) space (**Figure 4a** inset): For example, individual CSFs that are shifted more upward (peak-CS) are also shifted more rightward (peak-SF), and vice versa. In addition, bandwidth is correlated negatively with both peak-CS and peak-SF. Individual CSFs that are shifted more upward and rightward (higher peak-CS and peak-SF) are sharpened (narrower bandwidth). These significant correlations result in a more consistent organization of observers’ CSFs at the horizontal than at the other locations.

At the upper and lower vertical meridians (**Figures 4b-c**), there was a negative correlation between bandwidth and peak-CS/peak-SF, like at the horizontal meridian. However, at these vertical locations the peak-CS and the peak-SF were not correlated, indicating that individual CSFs do not necessarily vary along the diagonal direction. The fewer number of significant correlations results in less constraints on where different observers’ CSFs are located and how the attributes of their CSFs vary in the log(SF)-log(contrast sensitivity) space (**Figures 4b-c** insets).

For the covariation between cutoff-SF/AULCSF and all other attributes, see **Figure S6**.

## Discussion

The Contrast Sensitivity Function (CSF) is a building block of visual perception and an essential component of computational models of vision (Bradley, Abrams, & Geisler, 2014; Peli, 1996; Watson & Solomon, 1997). Thus, its variation with stimulus parameters has been widely assessed (DeValois & DeValois, 1990; Kelly, 1977; Graham, 1989; Watson, 2018), for example across eccentricity (Hilz & Cavonius, 1974; Jigo & Carrasco, 2020; Rovamo et al., 1978; Virsu & Rovamo, 1979). However, given the implicit or explicit assumption that CSF attributes generalize across iso-eccentric locations, scarce attention has been paid to variations of the CSF around the polar angle (Jigo et al., 2023; Kwak et al., 2024). Here, we conducted a detailed investigation of the key CSF attributes not only at a group level, line in our previous studies (Jigo et al., 2023; Kwak et al., 2024), but also at an individual level using covariance analysis. This analysis revealed critical differences within and across CSF attributes around the polar angle. Across locations, we fixed the CSF shape (bandwidth) for each individual and found that the CSF and its attributes -e.g., peak-CS and peak-SF-were enhanced and shifted to higher SFs at the horizontal than at the vertical meridian, and at the lower vertical than the upper vertical meridian. Moreover, each of the CSF attributes was highly correlated across polar angle locations. Within each location, the pattern in which individual CSFs vary was consistent, and more so at the horizontal than the vertical meridian.

With eccentricity, the peak-CS, peak-SF, and cutoff-SF of the CSF decrease, consistent with the drop in contrast sensitivity and spatial resolution in the periphery (Hilz & Cavonius, 1974; Jigo & Carrasco, 2020; Jigo et al., 2023; Rovamo et al., 1978; Virsu & Rovamo, 1979). As for bandwidth, results are more mixed. For example, Virsu & Rovamo (1979) reported that the attenuation in the low SF range becomes less pronounced at further eccentricities, indicating that the bandwidth increases from the fovea to the periphery. On the contrary, Jigo et al. (2023) found no changes in bandwidth between 2° and 6°. However, it should be noted that such results could be dictated by the choice of stimulus parameters (e.g., eccentricity, temporal frequency) known to affect the shape of the CSF (Graham, 1989; Watson, 2018), which differed across these studies. At the neurophysiological level, the bandwidth of V1 cells’ CSFs remain constant across eccentricity (De Valois, Albrecht, & Thorell, 1982).

Whether the bandwidth changes with eccentricity is not as clear as for other CSF attributes, and thus we compared models with fixed and varying bandwidths across polar angle locations for each individual. We found that CSFs at all meridians have the same *shape*: The most parsimonious model was the one in which the bandwidth varies across observers but is the same across locations for each observer. The shape of the CSF is related to the ratio of magnocellular and parvocellular cells in the LGN, which are sensitive to low and high SFs, respectively (Derrington & Lennie, 1984; Legge, 1978; Merigan & Eskin, 1986; Merigan, Katz, & Maunsell, 1991), and this ratio may be preserved at iso-eccentric locations around the visual field. If a template CSF with a fixed shape can account for various conditions (e.g., polar angle), the number of datapoints needed to be sampled in stimulus space would be greatly reduced. Some studies provide evidence in support of measuring the entire CSF (Bour & Apkarian, 1996; Hess & Howell, 1977; Rohaly & Owsley, 1993), whereas other studies claim redundancy of information in these procedures (Elliott & Whitaker, 1992; Pelli, Rubin, & Legge, 1986; Thurman, Davey, McCray, Paronian, & Seitz, 2016). Our findings suggest that, for each individual, CSFs at polar angle locations are of a fixed shape. This knowledge could greatly improve the efficiency in characterizing an observer’s CSF at various locations.

The group level sensitivity indicated by the key attributes was best for the horizontal meridian, followed by the lower vertical, and then the upper vertical meridian, consistent with previous studies (**Figure 2**; Jigo et al., 2023; Kwak et al., 2024). Therefore, although the shape of the CSF is preserved around polar angle, CSFs need to be shifted and scaled to account for these differences. At the 6° eccentricity tested in the present study, there were clear horizontal-upper and lower-upper asymmetries. The difference between the horizontal and the lower vertical meridian was present only for the peak-CS, but differences between the two locations might emerge for other attributes when tested at farther eccentricities where these asymmetries become more pronounced (Baldwin et al., 2012; Cameron et al., 2002; Carrasco et al., 2001; Himmelberg et al., 2020). These group-level behavioral effects are related to the higher density of cone photoreceptors and retinal ganglion cells along the horizontal than the vertical meridian, and of retinal ganglion cells at the lower than the upper vertical meridian (Curcio & Allen, 1990; Curcio, Sloan, Packer, Hendrickson, & Kalina, 1987; Kupers, Benson, Carrasco, & Winawer, 2022; Kupers, Carrasco, & Winawer, 2019; Song, Chui, Zhong, Elsner, & Burns, 2011; Webb & Kaas, 1976), as well as to the larger cortical surface area showing both asymmetries (Benson et al., 2021; Himmelberg et al., 2021, 2022).

In the current study, we also uncovered correlations at an individual level around the visual field. First, we found that each CSF attribute is highly correlated across polar angle locations (**Figure 3**), indicating that an individual’s CSF at one location is predictive of their CSFs at other polar angle locations. The CSFs are shifted and scaled by a similar amount for all observers across locations (e.g., a similar shifting or scaling factor across observers that explains how the CSF changes with locations); thus, the difference among individuals at a location is preserved around the visual field. At prima facie, this might seem inconsistent with the group-level CSF differences around polar angle. However, it is important to note that this correlation shows that an observer with higher horizontal sensitivity (e.g., peak-CS) than other observers also has a higher vertical meridian sensitivity than other observers; this is different from the *absolute* values of an observer’s sensitivity at these locations being similar. Taken together with the group-level CSF differences, we show that the absolute values of the CSF are *different* but *correlated* at an individual level around the visual field.

Moreover, the way in which individual CSFs vary is more consistent at the horizontal than at the vertical meridian (**Figure 4**). At the horizontal locations, observers whose contrast sensitivity is higher also peaks at higher SFs, which constrained individuals’ CSFs to vary diagonally in the log(SF)-log(contrast sensitivity) space: Individuals who are relatively more sensitive along the SF dimension are also more sensitive along the contrast dimension. Incorporating such relation between peak-CS and peak-SF into computational models of vision would enable a prediction of CSFs *across* individuals, which goes beyond predicting one observer’s CSFs at multiple locations. The HBM has been applied to predicting CSFs under different luminance conditions, across observers and for existing observers in unmeasured conditions (Lu, Zhao, Lesmes, & Dorr, 2022). A similar approach can be used for CSFs at different locations. Whether adding the constraint of the covariation between peak-CS and peak-SF into the HBM would increase its predictive power is an interesting question for future studies. However, at the vertical meridian, the covariation between peak-CS and peak-SF is less pronounced, highlighting that differences exist as a function of polar angle. Interestingly, De Valois et al (1982) found no correlation between the peak-CS and peak-SF of V1 cells’ CSFs. They reported that the recording loci varied from 0° to 5° eccentricity, but it is unclear where they recorded in terms of polar angle locations. Given that in our study, the correlation between the peak-CS and peak-SF across individuals was pronounced at the horizontal but not at the vertical meridian, it is plausible that collapsing measurements around the visual field obscured such effects in the previous study.

At both the horizontal and the vertical meridians, peak-SF/peak-CS were negatively correlated with bandwidth (note that although bandwidth was fixed across locations in our model, it varied with individuals). To explain these correlations, we speculate that there is a limit at the high SF range as well as on the total window of visibility (i.e., AULCSF) that the human visual system can process. If neurons can only process SFs up to a certain limit, the slope from the peak-CS to the cutoff-SF would be steeper, resulting in a narrower bandwidth for a CSF that peaks at a higher SF. Consistent with this speculation, there is neurophysiological evidence that the peak-SF and bandwidth of the CSF are negatively correlated across V1 cells (De Valois et al., 1982). Similarly, if there is a constraint on the area of the human window of visibility, a CSF peaking at a higher contrast sensitivity would be narrower.

In conclusion, the present study unveils important relations and differences among our window of visibility around the polar angle. Key attributes of the CSF vary with location, and the pattern in which CSFs differ across observers also varies between the horizontal and the vertical meridians. However, the shape of the CSF and individual variability in CSFs is preserved across locations. Altogether, these findings call for a closer investigation of factors that are similar and different as a function of polar angle location and of the underlying neural factors. Our study highlights that comprehensive models of vision should consider CSFs around polar angles, to better depict our window of visibility around the visual field.

## Methods

### Participants

Twenty-eight observers (14 males and 14 females, including author YK; ages 21-34) with normal or corrected-to-normal vision participated in the experiment. All participants except for the author were naïve to the experimental hypothesis. The experimental procedures were approved by the Institutional Review Board at New York University, and all participants provided informed consent. They were paid $12 per hour. All procedures were in agreement with the Declaration of Helsinki. Data of seven out of the twenty-eight participants are reported in a previous study (Kwak et al., 2024).

### Setup

Participants sat in a dark room with their head stabilized by a chin and forehead rest. All stimuli were generated and presented using MATLAB (MathWorks, Natick, MA, USA) and the Psychophysics Toolbox (Brainard, 1997; Pelli, 1997) on a gamma-linearized 20-inch ViewSonic G220fb CRT screen (Brea, CA, USA) at a viewing distance of 79 cm. The CRT screen had a resolution of 1,280 by 960 pixels and a refresh rate of 100 Hz. Gaze position was recorded using an EyeLink 1000 Desktop Mount eye tracker (SR Research, Osgoode, Ontario, Canada) at a sampling rate of 1 kHz. The Eyelink Toolbox was used for eyetracking with MATLAB and Psychophysics Toolbox (Cornelissen, Peters, & Palmer, 2002).

### Experimental Procedure

Participants performed an orientation discrimination task for stimuli varying in contrast and spatial frequency. **Figure 2a** shows the trial sequence. Each trial started with a fixation circle (0.175° radius) on a gray background (∼26 cd/m^2^) with a duration randomly jittered between 400ms and 600ms. Four placeholders indicated the locations of the upcoming stimuli, 6° left, right, above, and below fixation. Measurements at the left and right locations were combined for analysis, as it is well established that contrast sensitivity does not differ between these locations (Cameron et al., 2002). Each placeholder was composed of four corners (black lines, 0.2° length).

The trial began once a 300-ms stable fixation (eye coordinates within a 1.75° radius virtual circle centered on fixation) was detected. After detection of stable fixation, a non-informative cue -four black lines (each, 0.35° length) pointing to all locations-appeared. 140ms after the onset of the cue, a test Gabor grating (tilted ±45 relative to vertical, random phase, delimited by a raised cosine envelope with 2° radius) appeared for 30ms at one of the four locations. The spatial frequency and the contrast of the Gabor stimulus were determined by the quick CSF procedure on each trial, based on the maximum expected information gain (Lesmes et al., 2010). 450ms after stimulus offset, the location at which the test Gabor was presented was highlighted by increasing the width and length of the corresponding placeholders. The non-informative cue pointing to all directions changed to a response cue, which was a black line pointing to the test Gabor location. Participants performed an un-speeded, orientation discrimination task for the Gabor grating (by pressing left arrow key for -45°, right arrow key for +45°).

Participants were instructed to maintain fixation throughout the entire trial sequence. Gaze position was monitored online to ensure fixation within a 1.75° radius virtual circle from the central fixation until the response phase (fixation blocks) or until cue onset (saccade blocks). Trials in which gaze deviated from fixation were aborted and repeated at the end of each block.

To extract the CSF, we measured contrast thresholds at various spatial frequencies (**Figure 1c**). To place trials efficiently in the dynamic range of the CSF, we leveraged the qCSF (quick Contrast Sensitivity Function) procedure for data collection (Lesmes et al., 2010). Given participants’ performance in previous trials, this procedure selects stimulus contrast and spatial frequency to be tested on each trial, based on the maximum expected information gain (reduction in entropy), to further refine the CSF parameter estimates. The possible stimulus space was composed of 60 contrast levels from 0.001 to 1, and 12 spatial frequency levels from 0.5 to 16cpd, evenly spaced in log units.

### CSF analysis

#### Bayesian Inference

To characterize the CSFs for each location (upper, lower, horizontal), we used the Bayesian Inference Procedure (BIP, **Figure S1a**) and the Hierarchical Bayesian Model (HBM, **Figure S1b**) to fit the three-parameter CSF model to trial-by-trial data points (Zhao, Lesmes, Hou, et al., 2021). For both the BIP and the HBM, Bayesian inference was used to estimate the posterior distribution of the three CSF parameters - peak contrast sensitivity (peak-CS), peak spatial frequency (peak-SF), and bandwidth. Other key attributes, such as the cutoff spatial frequency (cutoff-SF) and area under the log CSF (AULCSF), were computed from the final estimate of the CSF. Because cutoff-SF and AULCSF are dependent on the three CSF parameters, we report results of these attributes in **Supporting Information**.

Contrast sensitivity at spatial frequency is modeled as a log-parabola function with parameters 0 = *(peakCS, peakSF, bandwidth)* Lesmes et al., 2010; Watson & Ahumada, 2005):

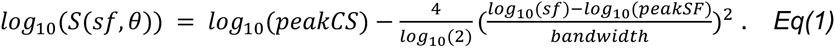

The probability of a correct response *(r* = 1) on a trial given the stimulus - spatial frequency *sf* and contrast *c* - is described as a psychometric function (**Figure 1a**):

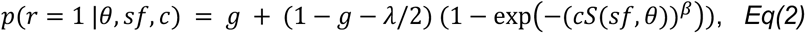

where *g* is the guessing rate (g = 0.5), *λ* is the lapse rate ( *λ* = 0.04; (Wichmann & Hill, 2001; Zhao, Lesmes, Hou, et al., 2021)), and *β* determines the slope of the psychometric function (/? = 2). The probability of making an incorrect response (r = 0) is:

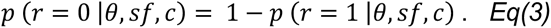

Equations 2 and 3 define the likelihood function, or the probability of a correct/incorrect response in a trial given the stimulus and the CSF parameters. To infer the CSF parameters given the experimental data, Bayes’ Rule is used to estimate the posterior distribution of the CSF parameters (6) based on the likelihood and the prior (*Eq(7)*) in the BIP procedure. Details on modeling the prior distributions can be found below (*Eq(8-10)*).

#### HBM: Three-level hierarchy

The BIP fits these parameters independently for each individual and condition (**Figure S1a**). Although the BIP has been proven to be a good estimate of the CSF, it may have overestimated the variance of each test because it scores each test independently with a uniform prior without considering potential relations of the parameters (Zhao, Lesmes, Dorr, & Lu, 2021). Therefore, we leveraged a hierarchical model which recently has been shown to reduce the uncertainties of the parameter estimates when fitting the CSF (Zhao, Lesmes, Dorr, et al., 2021; Zhao, Lesmes, Hou, et al., 2021). We used a three-level HBM with the same structure as in Zhao, Lesmes, Dorr, et al. (2021) (**Figure S1b**). In the current study, there were 28 individuals (participants; *I* = 28), 3 conditions (locations; *J* = 3), and all trials that each participant completed were combined into one test *(K* = 1). Note that in our previous work utilizing similar procedures (Kwak et al., 2024) we use a simpler three-level HBM which does not take into consideration the different conditions in the model structure (Zhao, Lesmes, Dorr, et al., 2021).

The HBM considers potential relations of the CSF parameters and hyperparameters within and across hierarchies. More specifically, it quantifies the joint distribution of the CSF parameters and hyperparameters at three hierarchies in a single model: test-(K), individual-(I), and population-level. The within-individual and cross-individual regularities across experimental conditions (1:/) are modeled as the covariance of the CSF parameters at the individual level *(p_7_j*) and covariance at the population level (£), respectively. The model incorporates conditional dependencies: CSF parameters at the test level are conditionally dependent on hyperparameters at the individual level, and the CSF hyperparameters at the individual level are conditionally dependent on those at the population level, Incorporating this knowledge into the model and decomposing the variability of the entire dataset into distributions at multiple hierarchies enabled us to reduce the variance of the test-level estimates and to obtain more precise estimates of the CSF parameters.

We fit the BIP and the HBM sequentially. As a first step, the BIP was fit to obtain the mean and standard deviation of each of the three CSF parameters across participants for each condition, using a uniform prior distribution. Next, these values were set as the prior for HBM (see *Eq(8-10)*).

At the population level of the HBM, the joint distribution of hyperparameter *η* across *J* experimental conditions was modeled as a mixture of three-dimensional Gaussian distributions *N* with mean / and covariance r, which have distributions p(/) and p(£):

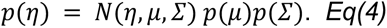

At the individual level (/), the joint distribution of hyperparameter *t_i,_*_1:*J*_ of individual *i* across all the 1: *J* experimental conditions was modeled as a mixture of three-dimensional Gaussian distributions *N* with mean *p_7_j* and covariance *φ_j_*, which have distributions *p*(*ρ_i,_*_1:*J*_ |*η*) and *p*(*φ_j_*); where *p*(*ρ_i,_*_1:*J*_ |*η*) denotes that the mean was conditioned on the population-level hyperparameter :

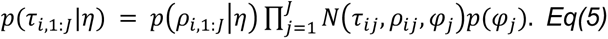

At the test level (*K*), *p*(*θ_ijk_*|*t_ij_*), the joint distribution of parameter *θ_ijk_* of individual *i*, condition *j,* and test *k,* was conditioned on hyperparameter *t_ij_* at the individual level.

The probability of obtaining the entire dataset *Y* was computed by probability multiplication:

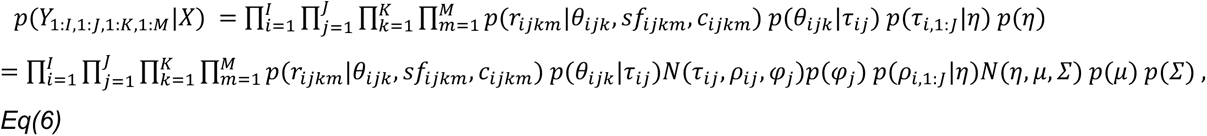

where *X =* (*θ*_1:*I*,1:*J*,1:*K*_,ρ_1:*I*,1:*J*_,*φ*_1:*J*_, *μ*, *Σ*) are all parameters and hyperparameters in the HBM, *r_ijkm_* is a response on a given trial, *sf_ijkm_* and *c_ijkm_* are spatial frequency and contrast of the stimulus on a given trial.

#### HBM: Computing the joint posterior distribution

Bayes rule (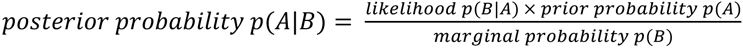) was used to compute the joint posterior distribution of all the parameters and hyperparameters in the HBM:

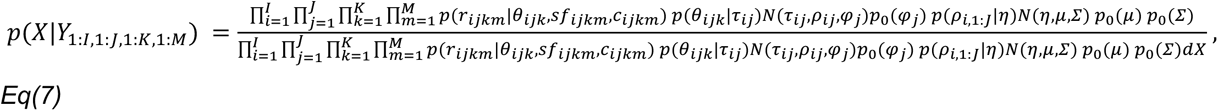

where the denominator is the probability of obtaining the entire dataset (*Eq(6)*).

We used the JAGS package (Plummer, 2003) in R (R Core Team, 2020) to evaluate the joint posterior distribution. JAGS generates representative samples of the joint posterior distribution of all the parameters and hyperparameters in the HBM via Markov Chain Monte Carlo (MCMC). We ran three parallel MCMC chains, each generating 2000 samples, resulting in a total of 6000 samples. Steps in the burn-in and adaptation phases -20000 and 500000 steps respectively-were discarded and excluded from the analysis because the initial part of the random walk process is largely determined by random starting values.

### Prior distributions in the HBM

For fitting the HBM, we start with prior distributions of *μ* (population mean), *Σ*^-1^ (inverse of population level covariance matrix), and *φ^-1^* (inverse of individual level covariance matrix), which are *p*_0_(*μ*), *p*_0_(*Σ*^-1^), and *p*_0_(*φ^-1^*) respectively.

For each of the CSF parameter, the prior distribution of *μ* is a uniform distribution:

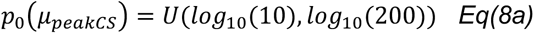

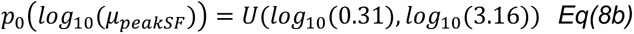

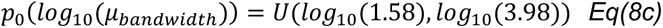

The prior distributions of the precision matrices (the inverse of covariance matrices *X* and p) are modeled as Wishart distributions. *W(Y,v)* denotes a Wishart distribution with expected precision matrix *Y* and degrees of freedom *v (v* = 4) *Σ_BIP_* and *φ_BIP_* are population-level and individual-level covariance matrices obtained from the BIP fit:

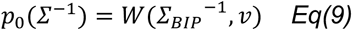

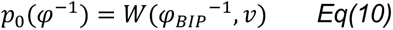

### Model comparison

To directly test whether the shape of the CSF differs around the polar angle, we compared two models - one in which bandwidth was fixed across locations (fixed bandwidth model) for each observer, and one in which bandwidth was free to vary across locations (varying bandwidth model) for each observer. In the fixed bandwidth model, there were 7 parameters per participant: 2 parameters (peak-CS, peak-SF) x 3 locations + 1 parameter (bandwidth). In the varying bandwidth model, there were 9 parameters per participant: 3 parameters (peak-CS, peak-SF, bandwidth) x 3 locations. Here, we report results from the fixed bandwidth model, which is more parsimonious. That the current study used a fixed bandwidth model explains why the pattern in peak-SF (**Figure 2c**) and bandwidth across polar angle is not identical to that in prior work using a varying bandwidth model (Jigo et al., 2023; Kwak et al., 2024).

### Covariance matrix

The 15-by-15 covariance matrix across all attributes and locations (5 attributes x 3 locations = 15 combinations) was obtained by computing the covariance of a 28-by-15 (28 participants, 15 combinations) matrix. Shrinkage estimates of the covariance matrices were computed with package “corpcor” in R. The correlation matrix of the covariance matrix was obtained with the MATLAB function “corrcov”, which returns the Pearson correlation coefficients. The confidence intervals for the correlation coefficients were computed from the distribution of the MCMC samples (**Figure S7**). Note that only the 9-by-9 matrix (3 attributes x 3 locations = 9 combinations) is presented in the main text, and other attributes are reported in **Supporting Information**.

We also report the variation across individuals (coefficient of variation) for each CSF attribute, to compare the variation for each attribute which has different units (**Table S1**).

### Statistical analysis

Convergence of the HBM parameters was determined based on the Gelman and Rubin’s diagnostic rule (Gelman & Rubin, 1992). Each parameter and hyperparameter was considered to have “converged” when the variance of the samples across the MCMC chains divided by the variance of the samples within each chain was smaller than 1.05. In the current study, all parameters of the HBM converged.

To obtain final estimates of the CSF parameters in the HBM -peak-CS, peak-SF, and bandwidth-, parameter estimates of the 6000 samples were averaged for each participant and location. The final CSFs for each participant and location were obtained by inputting the final parameter estimates to *Eq (1)*. From these final CSFs, cutoff-SF and AULCSF were computed. These key CSF attribute values were used for statistical testing.

*P* values for the average CSF key attributes are based on permutation testing over 1000 iterations with shuffled data: the proportions of *F* score or *t* scores in the permuted null distribution greater than or equal to the metric computed using intact data. For covariation of attributes at each location and covariation of locations for each attribute, we compared the distribution of *p* values of correlation coefficients (obtained from 1000 MCMC samples) against an arbitrary cutoff value of 0.05. Note that the minimum *p* value achievable with these procedures is 0.001. All *P* values were FDR corrected for multiple comparisons, when applicable (Benjamini & Hochberg, 1995).

## Acknowledgements

We thank Yukai Zhao, Jonathan Winawer, as well as Rania Ezzo, Aysun Duyar, and other members of the Carrasco Lab for helpful discussion and comments. This research was supported US NIH National Eye Institute R01-EY027401 to MC.

## Supporting Information

**Figure S1.**
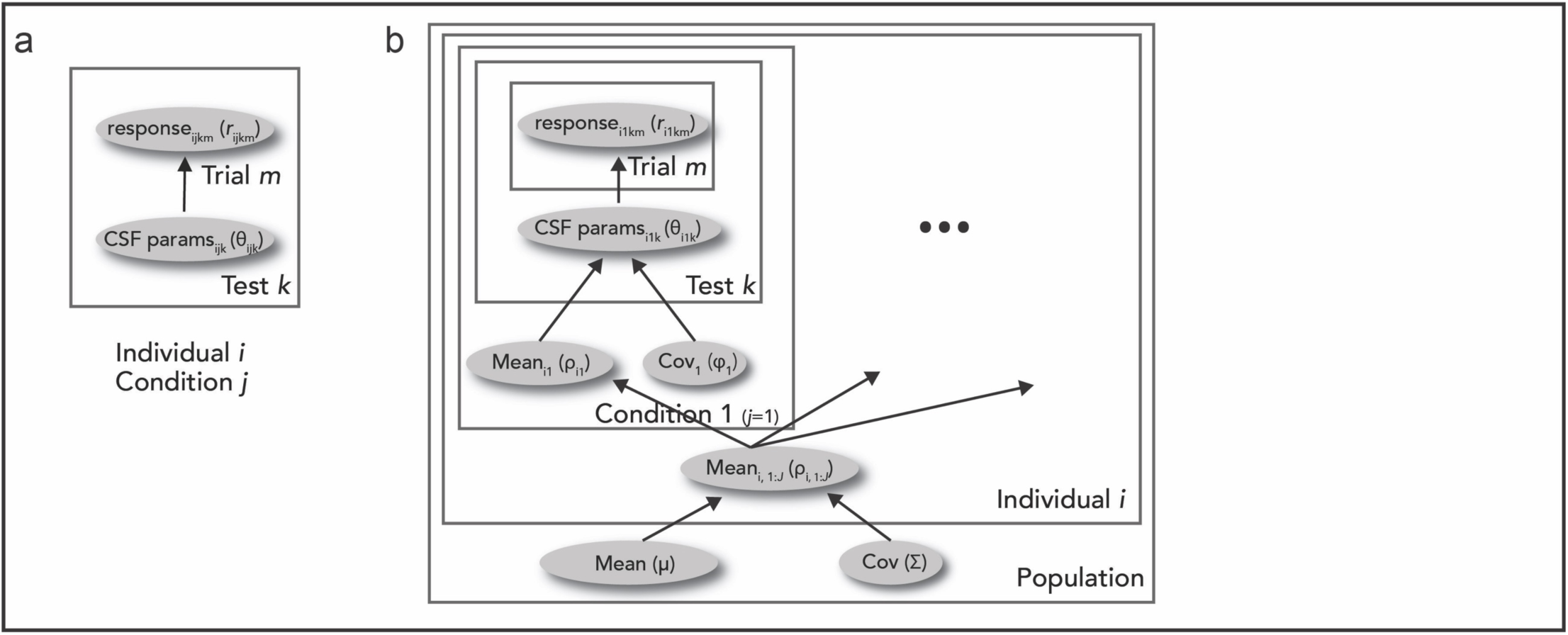
CSF model fitting. **(a)** Schematic representation of the BIP (Bayesian Inference Procedure). The first stage of the data analysis consists of fitting trial-by-trial data points with the BIP to estimate the CSF parameters - peak-CS, peak-SF, and bandwidth. The BIP computes the posterior distribution of the parameters for each test independently. **(b)** Schematic representation of the HBM (Hierarchical Bayesian Model). In the second stage of the data analysis, the BIP outputs are used as the mean and covariance of the prior distributions of the CSF hyperparameters in the HBM. We used a three-level hierarchical model to incorporate potential relations in the CSF parameters across individuals and tests. For details on the BIP and the HBM, see **Methods**.

**Figure S2.**
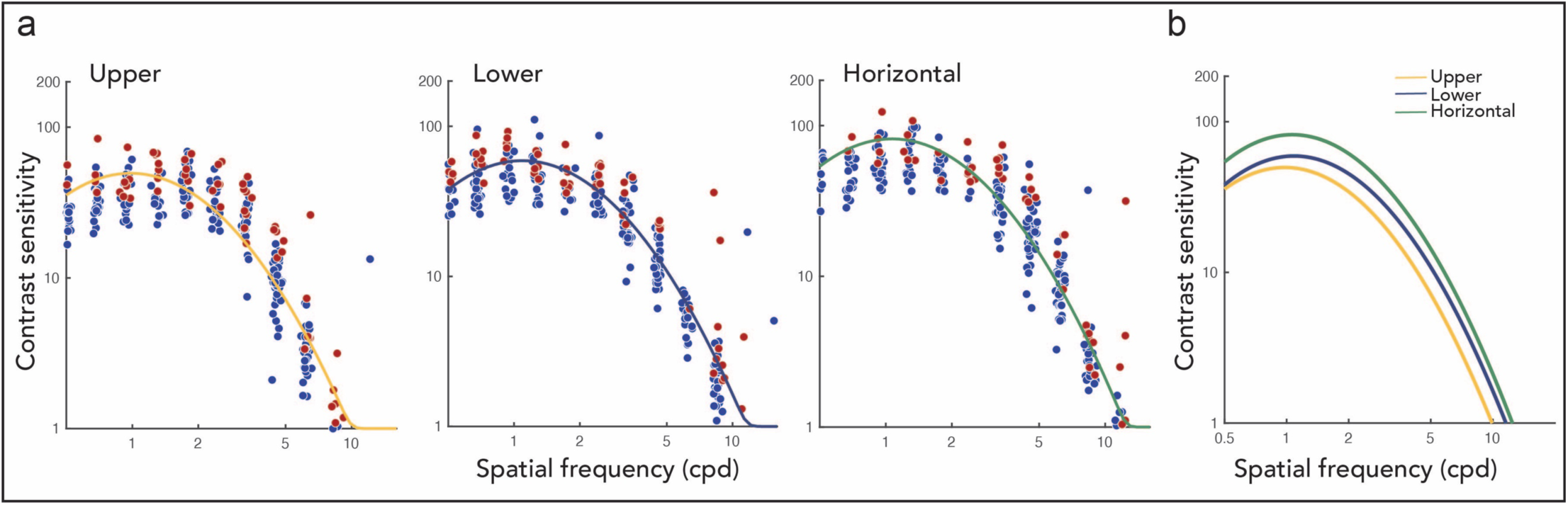
Data from an example observer. **(a)** Trial-by-trial datapoints (red for incorrect trials, blue for correct trials) and fitted CSFs for the upper, lower, and the horizontal meridians. **(b)** CSFs across locations are overlayed.

**Figure S3.**
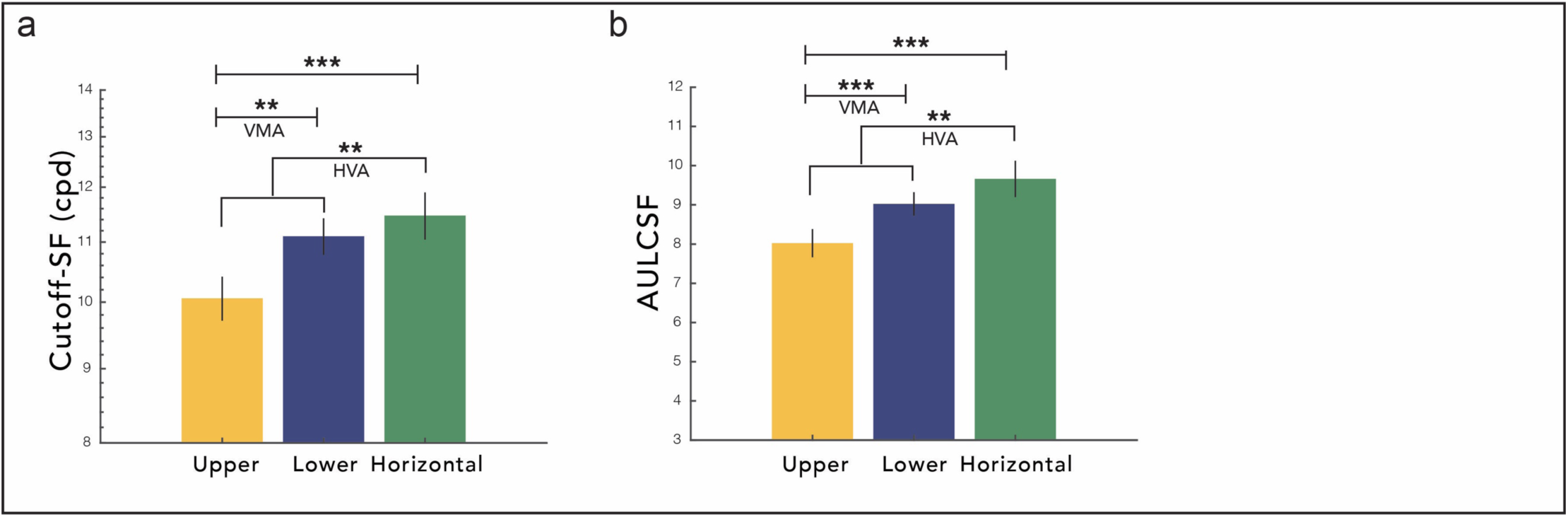
Polar angle sensitivity differences, **(a) Cutoff spatial frequency, (b) Area under the log CSF.** *\*\*\*p* < 0.001, *\*\*p* < 0.01. Error bars are ±1SEM. HVA and VMA denote Horizontal-Vertical Anisotropy and Vertical Meridian Asymmetries, respectively.

**Figure S4.**
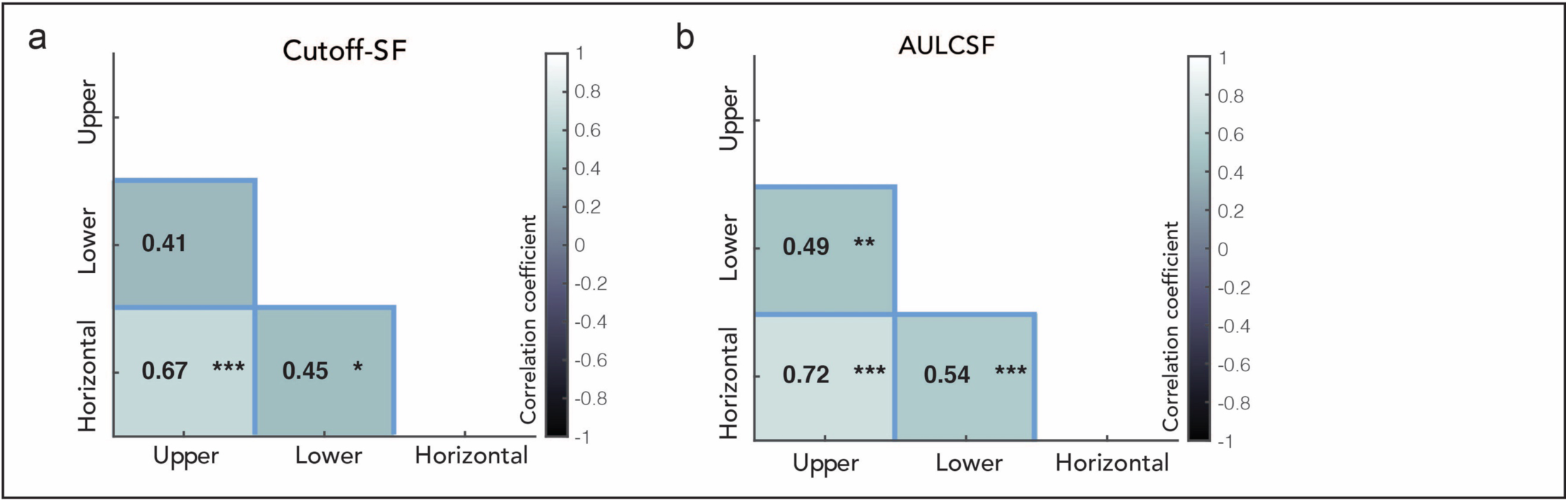
Covariation of each CSF attribute across polar angle locations, **(a) Cutoff spatial frequency, (b) Area under the log CSF.** Confidence intervals are in **Figure S7.** *\*\*\*p* < 0.001,**p < 0.01, *\*p* < 0.05.

**Figure S5.**
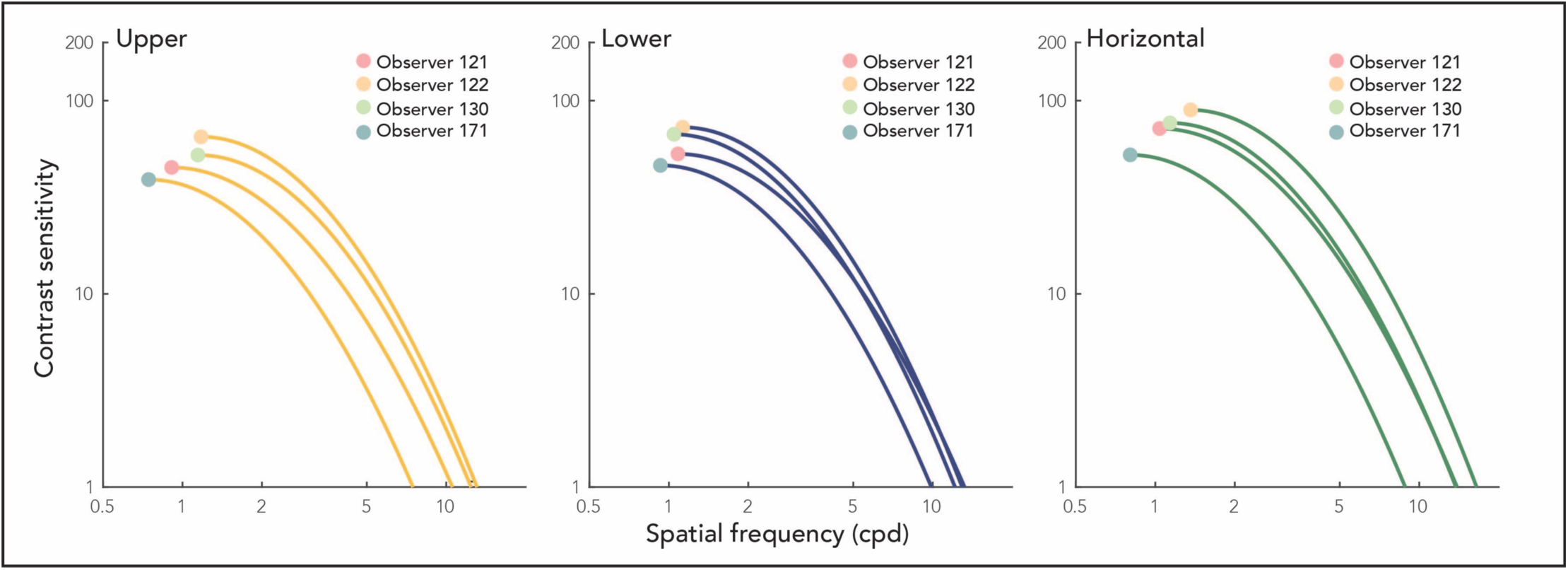
Data from four example observers at each location. Each circled dot indicates the peak-CS and peak-SF of each observer. Note that across locations, observers with higher peak-CS and peak-SF at one location tend to have higher peak-CS and peak-SF at other locations, indicating that CSF attributes are highly correlated across polar angle (Figure 3). Moreover, at the horizontal meridian, observers’ CSFs are shifted consistently along the diagonal direction in the SF-contrast space (Figure 4), where the peak-CS and peak-SF fall closely along a diagonal line. Only the part of the CSF corresponding to spatial frequencies above the peak-SF is shown for a clearer illustration of peak-CS and peak-SF.

**Figure S6.**
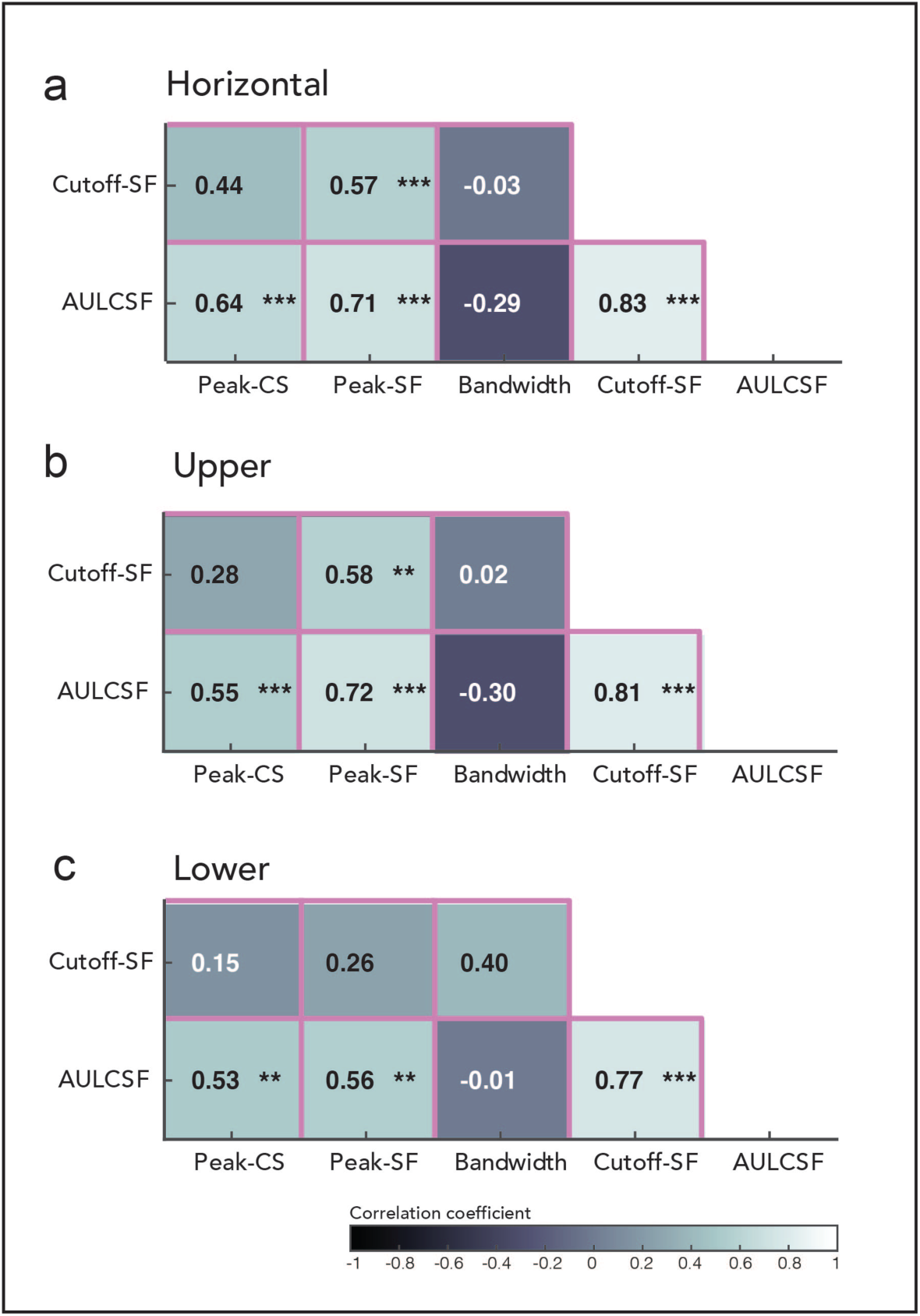
Covariance of CSF attributes within each location (pink cells in Figure 1d-e). Correlation between pairs of attributes at the **(a)** Horizontal, **(b)** Upper vertical, and **(c)** Lower vertical meridian. The color of each cell corresponds to the strength of correlation. Each cell corresponds to a pink cell in Figure 1d. Confidence intervals are in **Figure S7**. \*\*\**p* < 0.001, \*\**p* < 0.01. Black and white letters are for visibility.

**Figure S7.**
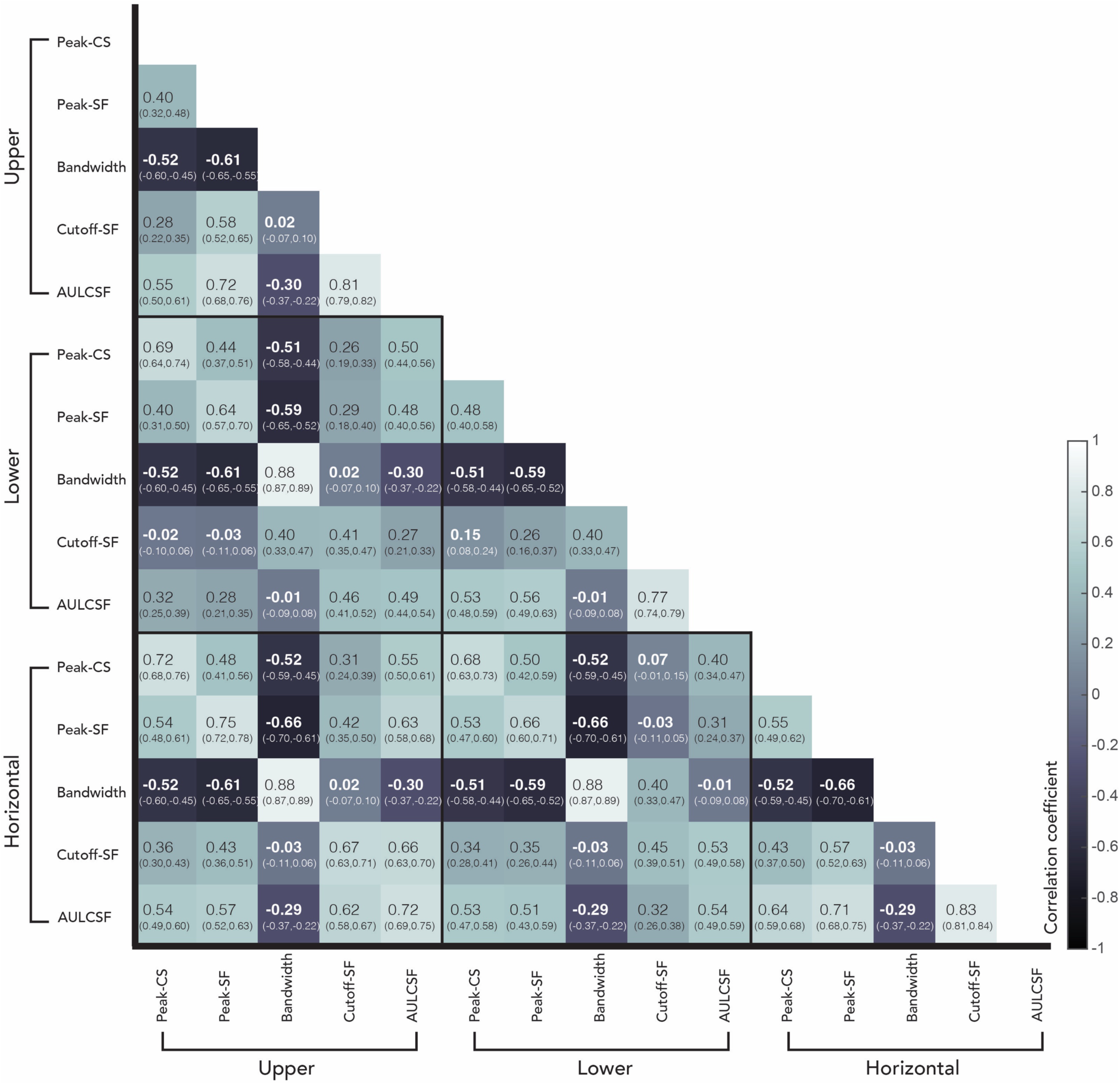
Correlation coefficients and confidence intervals. Correlation coefficients are in the center of each cell, and confidence intervals are in the parentheses below. The distribution of MCMC samples (**Methods**) was used to compute the 68% confidence interval. Black and white letters are for visibility.

**Table S1.** Coefficient of variation (CoV). For each attribute at each location, we computed the CoV (ratio of the standard deviation to the mean), which is useful for comparing the variation across individuals for each CSF attribute which differs in units.

|  | Upper | Lower | Horizontal |
| --- | --- | --- | --- |
| Peak-CS | 0.30 | 0.25 | 0.25 |
| Peak-SF | 0.23 | 0.14 | 0.24 |
| Bandwidth | 0.18 | 0.14 | 0.20 |
| Cutoff-SF | 0.19 | 0.15 | 0.20 |
| AULCSF | 0.24 | 0.18 | 0.25 |

